# Improving the RNA velocity approach using long-read single cell sequencing

**DOI:** 10.1101/2022.05.02.490352

**Authors:** Chen Zhang, Weitian Chen, Yitong Fang, Zhichao Chen, Yeming Xie, Wenfang Chen, Zhe Xie, Mei Guo, Juan Wang, Chen Tan, Hongqi Wang, Chong Tang

**Affiliations:** BGI Shenzhen, China, 518000; College of Life Sciences, University of Chinese Academy of Sciences, Beijing, 100049, China; Department of Biology, Cell Biology and Physiology, University of Copenhagen 13, 2100 Copenhagen, Denmark

**Keywords:** single-cell, ONT full-length sequencing, three-barcode technology, single cell maps, RNA velocity

## Abstract

The concept of RNA velocity has been recently developed that allowed to look at the otherwise static single-cell RNA sequencing data in a dynamic way, which permitted inferences about cell fates. However, the more precise parameters, such as the number of exons/introns, can also be determined using long-read methods. Comparing the numbers of exons and introns allows including more genes for downstream velocity analysis and resolves the precise cell fate. The recently developed concept of “RNA velocity” concerns with dynamic changes in mRNA expression and complements single-cell RNA sequencing (scRNA-seq) data, which are static snapshots of a certain cell state taken at a given time point^1^. RNA velocity measures the change in mRNA abundance by differentiating the newly transcribed unspliced pre-mRNAs from mature spliced mRNAs. The rapidly developing long-read sequencing technology lends itself for RNA velocity analysis of scRNA-seq data, which was previously performed primarily using second-generation sequencing.

---

Third-generation sequencing is limited by the low throughput and lack of accuracy. Our groups developed two high-throughput third-generation sequencing methods for different single cell platforms. The high-throughput single-cell full-length isoform sequencing method (HIT-scISOseq) ligated multiple reads (10X Genomics) into a single molecule, which quadrupled the throughput on the Pacific BioSciences (PacBio) platform^2^. In current research, we have developed a three-barcode technology, high-throughput single-cell Oxford Nanopore full-length RNA sequencing (HISOFA-seq), which is accessible to most researchers. The ultralong three barcode is discriminated by the nanopore technology with 70% de-barcoding efficiency, generating 70 million reads (BD Rhapsody) per lane (Extended Data Fig. 1).

HIT-scISOseq/HISOFA-seq enables measurement of the key RNA velocity parameters, counts of unspliced and spliced mRNAs, which are mainly determined by the presence or absence of introns^3^. Given that second-generation sequencing has short read length, attributing all short reads without introns to spliced mRNA may be inaccurate. Although spliced and unspliced mRNA counts strongly correlate in both the second-generation (NGS) and third-generation (PacBio) scRNA-seq data, the slopes of the relationships between spliced and unspliced counts significantly differed in these two datasets (Fig. 1b). In the NGS data, the fraction of unspliced counts was 8.0%, whereas that in the PacBio data comprised 24.1% (Fig. 1b). The apparently underestimated ratio of unspliced counts may complicate RNA velocity analysis. We explored this hypothesis in the well characterized biological model of spermatogenic development^4^ (Extended Data Figs. 2, 3). The original framework of RNA velocity was applied to obtain the vector fields projected onto the t-distributed stochastic neighbor embedding (tSNE) plot with observed and extrapolated velocity values of sperm cells from the 10x scRNA-seq data and HIT-scISOseq data, respectively, which showed their differences in the tSNE dimension of cell clustering and cell fate indicated by vector fields (Fig. 1f; Extended Data Fig. 4a).

**Figure 1.**
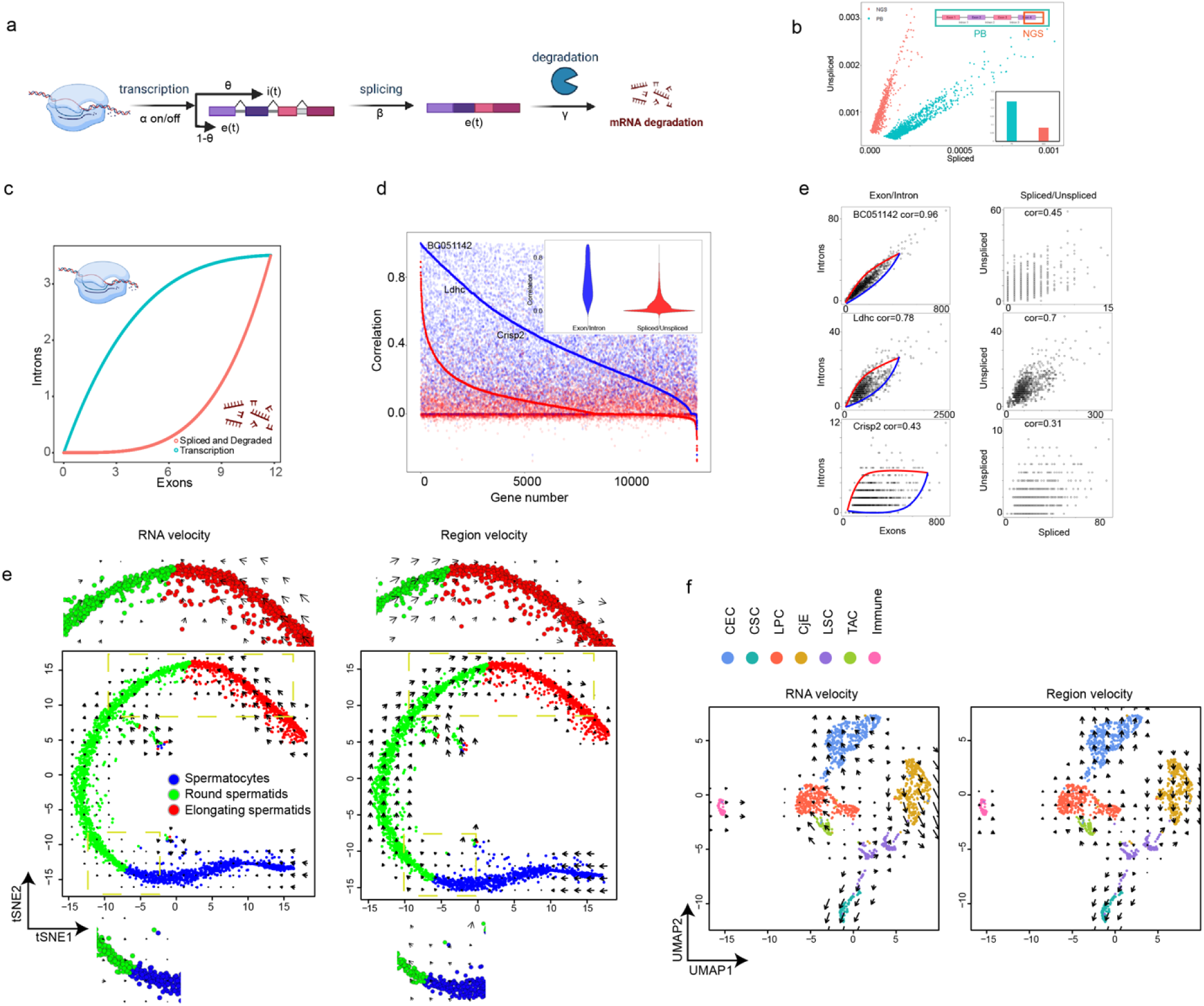
Principles and performance of Region velocity. (a) The change of exons and introns in the model of RNA transcription dynamics, capturing ratio of introns (θ), transcription (α), splicing (β), degradation (γ) rates. (b) The difference of spliced counts and unspliced counts between NGS and Pacbio data including their distribution and ratio. Blue indicates Pacbio data while red indicates NGS data. The histogram is the ratio of introns in NGS and PB data. (c) The simulation of exons and introns using Region velocity. The time of exons and introns of simulation is predicted from 0 to four times of transcription change time (ts) using parameters from Region velocity. The exons and introns are determined by the analytical formula in which the parameters are obtained from steady state models. Each point means each cell. Blue indicates cells are in transcription process and red indicates cells are in splicing or degradation process. (d) The difference of correlation between exons : introns and spliced counts : unspliced counts. Blue indicates the correlation between exons and introns while red indicates the correlation between spliced counts and unspliced counts. The lines were drawn based on the order of correlation values from highest to lowest. (e) The specific genes further demonstrated the difference in figure 1d. Three genes in the left represented three quantile of their correlation values of exons and introns and in the right the corresponding genes of unspliced counts and spliced counts are chosen. (f) Velocity filed projected onto tSNE plot of RNA velocity and Region velocity from full-scale scRNA data of mouse spermary cells (n=3,001 cells) using HIT-scISOseq. Arrows means the average speed on a defined grid (number of grids=20). Blue points represent spermatocytes, green points represent round spermatids and red points represent elongating spermatids (Cell population shown in Extended Data Fig. 3). (g) Velocity filed projected onto tSNE plot of RNA velocity and Region velocity from full-scale scRNA data of cynomolgus monkeys’ limbal cells (n=1,417 cells) using HIT-scISOseq. Cell population, CEC, Corneal epithelial cells; LPC, Limbal progenitor cells; CjE, Conjunctival epithelium; LSC, Limbal stem cells; CSC, Corneal stromal cells; TAC, Transient amplifying cells; Immune, T cells or immune cells. See also Extended Data Fig. 8.

Besides spliced and unspliced counts, HIT-scISOseq/HISOFA-seq generate more detailed information, which can be used to further develop the RNA velocity concept. HIT-scISOseq/HISOFA-seq detects full-scale mRNAs (from nascent to degrading), the lengths of which are suitable for RNA velocity calculations. According to RNA velocity modeling dynamics, mRNA length also varies over time in the process of transcription, splicing, and degradation. The vector fields projected onto the tSNE plot of the RNA velocity length model with mRNA length can be drawn, which is similar to the output of the original model using unspliced and spliced counts, except the vector fields are larger (Fig. 1f; Extended Data Fig. 4a, b). The similarity may be caused by the relatively low influence of mRNA length compared to that of RNA abundance. Both models did not produce expected sequence of spermiogenesis biological development.

As HIT-scISOseq/HISOFA-seq generates a substantial amount of data, we extended the RNA velocity model from the original framework using more robust parameters. HIT-scISOseq/HISOFA-seq also determines the numbers of introns and exons of each gene or isoform, which can substitute the spliced/unspliced counts and generate a more precise steady-state model, as a wide range of intron counts might produce different velocity estimates, whereas unspliced counts attribute genes with a different number of introns to the same unspliced count (Extended Data Fig. 5a, b). During transcription, the numbers of both introns and exons in nascent mRNA increase, whereas splicing and degradation decrease the numbers of introns and exons, respectively (Fig. 1a). Therefore, the determination model of this basic kinetic reaction is described as follows for each gene independently:

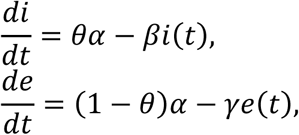

where *θ* is the proportion of introns in the transcription process; *e* and *i* are the numbers of exons and introns; *β* and *γ* are the splicing and degradation rates. As the state of the cell in the whole process is deduced by the time derivatives of the numbers of intragenic and transcribed regions, we termed this state change process as “Region Velocity”.

Region velocity is primarily observed through the spindle-shaped relationship between the number of exons and introns in different genes, representing a steady-state model of the original RNA velocity parameter, and their correlation level varies in different genes^3^. Data simulation showed that the levels of introns above or below the identity line may indicate the active state of transcription or splicing and degradation, respectively (Fig. 1c). To infer the unknown parameters (α, β, γ, and θ), we built a basic steady-state model and exploited the accurate inference by the dynamic model (see online Methods). Compared to RNA velocity, region velocity is more stable and better suited for steady-state modeling. Mathematically, the higher correlation between spliced/exon counts and unspliced/intron counts in different cells means shorter distance to the steady-state line (fitted line), indicating more reasonable projection of cells to the state of the next time point obtained using steady-state model parameters. The coefficients of correlation between unspliced and spliced counts in different cells were much lower than those of intron/exon counts, and 55.9% of genes had the intron/exon correlation coefficient >0.4 (Fig. 1d, e). Comparing the numbers of exons and introns allowed including more genes for downstream velocity analysis, as abundant zero values of spliced counts prevented calculation of their correlation with unspliced counts (Extended Data Fig. 5c, d). The dynamic model of RNA velocity using the expectation–maximization (EM) algorithm was designed to optimize steady-state modeling, because some genes have a short steady-state stage^5^. However, as more genes closer to the steady-state condition could be observed using region velocity calculation from exon and intron counts, the EM algorithm was not as fundamental for region velocity as for RNA velocity (Extended Data Fig. 6). In contrast, a steady-state model of region velocity had a satisfactory performance. Figure 1f shows vector fields projected onto the tSNE embedding plot of the steady-state model of the HIT-scISOseq data from sperm cells. Compared to RNA velocity calculations based on mRNA abundance or length, the direction of flow of region velocity was more aligned with expected spermatogenic waves: a streamlined flow from spermatocytes to elongating spermatids was evident^4^. As could be inferred from the length of the velocity field, RNA velocity accelerated at the beginning and decelerated at the end of the process of transformation of spermatocytes into round spermatids. This is consistent with increased RNA transcription at the beginning and during long periods of the two-step meiosis^4^. In spermatids cluster, the speed was increased at the round stage and decreased between the round stage and the start of elongation, suggesting that the elongation period lasted longer. Therefore, region velocity calculations indicated that meiosis and elongation are the speed bottlenecks of spermatogenesis, and their effects were similar based on the size of velocity field lengths. In addition, HIT-scISOseq data from the injured corneal epitheliums of cynomolgus monkeys were also analyzed using region velocity (Extended Data Fig. 8). Vector fields projected onto UMAP embedding plot for this dataset are illustrated in Fig. 1g. The direction flows of RNA velocity and region velocity both demonstrated transformation of limbal stem cells (LSCs) into terminally differentiated cells, including corneal epithelial and stroma cells^6^. However, region velocity identified LSCs as the differentiation center more precisely than RNA velocity. The arrows in the center of LSCs were clearly directed to the periphery, from a point to circles. Additionally, RNA velocity indicated conjunctival epithelium and corneal epithelial cells as origin, which contradicted known biological characteristics (Fig. 1g). Therefore, region velocity had better performance than RNA velocity in the monkey limbal wound model.

The third-generation HISOFA-seq platform is based on nanopore sequencing, whereas HIT-scISOseq uses PacBio sequencing. Region velocity is more applicable to HISOFA-seq data than RNA velocity. As nanopore sequencing has a higher error rate, gene counts obtained after isoform clustering yielded better performance for region velocity calculations than original gene counts. The vector fields projected onto the tSNE embedding plot indicated a consistent streamlined flow of spermatogenic waves in region velocity (Extended Data Fig. 7).

Therefore, region velocity is a multi-platform and multi-model parameter to project cell state, which is based on long-read scRNA-seq. In the future, HIT-scISOseq/HISOFA-seq may provide more data to discover new kinetic models of RNA dynamics, enabling better cell fate prediction for various species and different cell types.

## Method

The experimental protocols (HISOFA-seq) could be found on https://www.protocols.io/private/CE43BD78C53111EC97780A58A9FEAC02. The HIT-scISOseq protocol could be found on https://www.protocols.io/private/7472E845C45C11EC97780A58A9FEAC02.

The bioinformatic script could be found on https://www.protocols.io/private/17764990CA1611EC89580A58A9FEAC02 and available data could be found on https://github.com/Dekayzc/Regionvelocity/tree/main/data.

### Demultiplexing barcodes

#### Reference construction

BD cell label is roughly composed of five sections, including three separated barcodes of 9bp and two known linkers. Each barcode is randomly selected from a pool of 96 pre-defined sequences which allowed us labeling up to 96^3^ cells. The minimum Hamming distance of three barcodes was 4 bp and the mean distance was 6.6 bp. If taking insertion, deletion and substitution events into account, their minimum Levenshtein distance was 2 bp and the mean was 5.7 bp. In comparison to 10X barcode, BD barcode takes the advantage of its longer design and therefore has a greater edit distance.

We constructed the cell label reference by generating 96^3^ combinations of 9bp barcodes joined by two known linker sequences, so at this stage, the reference should begin with the first barcode and end with the third barcode. However, problems would arise with this type of reference because its flanking regions of may be clipped off during alignment. In order to avoid this happening, we elongated the reference with more sequences to act as the anchor, including binding site, UMI and oligo-dTs. Therefore, the final version of reference had a length of 98 bases, and we added more Ns to both ends to ensure the reference is longer than reads.

#### First round of demultiplexing

We took a subset of sequencing data and had a rough inspection of the library. Based on the positions of oligo dT, around 47% of the reads were classified as forward strands and 40% reads were reverse strands. Nearly 60% reads could be fully aligned to 5’ and 3’ binding site, indicating most of the reads possessed a complete library structure.

In order to find the most appropriate method to assign barcodes, we examined the performance of various aligners including minimap2, bwa-mem, bowtie2 and BLASTN. Although minimap2 was usually recommended for long read alignment, only less than 25% reads could be successfully demultiplexed. Similarly, many reads with the correct cell label were either wrongly assigned or unmapped in bwa-mem algorithm. As such, they were less recommended to use in this case where the length of query sequence was much longer than the reference. In comparison, the local alignment mode of bowtie2 and BLASTN outperformed other tools by having a mapping rate of 57.87% and 76.92%, respectively. BLASTN alignment generated less gaps, mismatches and had a longer mean aligned length than bowtie2, suggesting a better accuracy. However, bowtie2 was superior due to its ultrafast and memory-saving features. The speed of demultiplexing was further boosted when the read was trimmed into 300bp based on theoretical positions of barcodes. For example, bowtie2 and BLASTN could demultiplex 400,000 and 4,000 reads respectively under the same clock time. All these results suggested that bowtie2 was much more efficient than BLASTN in this case, thus the local alignment mode of bowtie2 became the optimal choice for the first round of demultiplexing.

#### Validation and optimization of the first round of demultiplexing

Our demultiplexing strategy was further validated using simulated data, which was created for the purpose of mimicking the real condition by introducing sequencing errors and PCR errors [1]. As table xxx shows, the true positive rate (TPR), false positive rate (FPR) and false negative rate (FNR) of BLASTN were 72.5%, 6.4% and 21.1%, respectively. The high precision value (0.919) also indicated the result was reliable enough for downstream analysis. In comparison, bowtie2 had a much worse precision of 0.719 due to its high FPR (27.9%), which would affect the accuracy of our demultiplexing result and subsequent analyses. Therefore, we carried out more attempts on improving and optimizing on bowtie2 parameters in order to reduce its false positive to the minimal level. The optimized alignment had a TPR of 70.7% and its FPR was dramatically reduced to 2.4%, which was even lower than BLASTN. Due to its high TPR and low FPR, bowtie2 with optimized parameters turned out to be the best strategy for the first round of demultiplexing.

#### Second round of demultiplexing (Optional)

During the first round of demultiplexing, the procedure of trimming the reads into 300bp was strict and harsh, and those reads with barcode outside the proposed positions might be lost. In addition, it could not tolerate the cases when three barcodes were linked by defective linkers. Therefore, it was necessary to rescue those reads with another round of demultiplexing.

In contrast to the previous step where reads were aligned to the cell label, we did not use cell label as the reference for the second step. Three barcodes were individually aligned to the reference, which was constructed by all unassigned reads from the first round of demultiplexing. If a read was aligned to all three barcodes and they were separated by an interval of similar length to the linker sequence, it will be assigned to the corresponding cell label. Using the test dataset, this procedure could further rescue 13.7% reads, which increased the demultiplexing rate from 57.87% into 71.58%. All unmapped reads will be discarded for downstream analysis.

#### Single-cell short read analysis

single-cell expression matrix was generated by standard 10X Genomics CellRanger pipeline (version 3.1.0).

### Single-cell long read analysis

#### Pacbio Single-cell isoform sequencing analysis

Generation of Circular Consensus Sequencing Reads, generation of Single Cell Full-Length Non-Concatemer (FLNC) Reads, genome alignment of FLNC reads, Cell Barcode correction and UMI correction, collapsing redundant isoforms, were performed according to HIT-scISOseq (Zheng et al., 2020).

#### Nanopore Single-cell isoform sequencing analysis

##### Genome alignment

The Full length of reads from the previous steps are mapped to a supplied reference genome(GCF_000364345.1_Macaca_fascicularis_5.0, mm10 (GENCODE vM23/Ensembl 98))using minimap2(v2.17-r954-dirty)(H. Li, 2018) with the following parameters: -ax splice -uf --secondary=no -C5. samtools (v1.9) (Heng Li et al., 2009) was used to compress sam files into bam, sort and index.

##### Generate gene-barcode matrix

spliced_bam2gff (https://github.com/nanoporetech/spliced_bam2gff) with default parameter was used to convert spliced bam alignments into GFF2 format, The GFF2 file was then compared with the reference comments using the gffCompare(v0.11.6) (Pertea & Pertea, 2020) tool. Besides the exon related class code :” = c k m n j e o “, intron related class_code “i,y” was selected for further analysis. Then gene-barcode matrix was generated by Barcode file, UMI file read and gene mapping file through our custom script. Collapsing redundant isoforms

We used cDNA_Cupcake python scripts (https://github.com/Magdoll/cDNA_Cupcake) python script collapse_isoforms_by_sam.py to collapse redundant isoforms, the parameters were set: as -c 0.95 -i 0.95 --max_fuzzy_junction 5 --max_5_diff 1000 --max_3_diff 30.

##### Generate exon and intron matrices

In order to know whether reads are from spliced transcripts or unspliced transcripts, we need to see if reads contain intronic sequences.

For short reads RNA single cell sequencing, exon and intron count matrices were peformed by velocyto (v0.6) (La Manno et al., 2018).

For long reads RNA single cell sequencing, the known genomic exon and intron coordinates were extracted from the GTF annotation file, and the overlapping coordinates of exon and intron coordinates were merged using BedTools merge respectively. Then we extracted the coordinates of splice alignment of observation data and compared them with known exon and intron annotation through BedTools intersect. Some researchers believe that the intron length is greater than 50bp (Piovesan et al., 2015), while others believe that the minimum intron length is 20bp (Jon et al., 2008). Therefore, an intersection with known intron with a length greater than 20bp was considered as an intron, where the count of intron with the length of 20bp-50bp accounts for only 0.5% of the known genomic intron annotation file.

##### Generate spliced and unspliced matrices

A long read with intronic sequences were considered as unspliced transcripts and cell-gene-unspliced/unspliced matrices was generated by our custom python script. cell-calling using gene-barcode matrix R package Matrix (v1.4) was used to load data as sparse Matrix, and barcodeRanks from DropletUtils(v1.2.2) (Lun et al., 2019) was used to calculate the Inflection and Knee of barcode rank and UMI distribution plot. And then the identification of cells from empty droplets was performed by emptyDrops function (Lun et al., 2019), threshold of gene counts (less than 20)for barcodes were specified as background. FDR (0.01)for testing whether a barcode is a empty cell.

Cells calling according to the number of UMIs associated with each barcode performed by defaultDrops function (Lun et al., 2019). Finally, Estimated Number of Cells, Total Genes Detected, Mean Genes per Cell, Median Genes per Cell, Mean UMI Counts per Cell, Median UMI Counts per Cell, Mean Reads per Cell, Median Reads per Cell and Fraction Reads in Cells, as implemented in CellRanger, were calculated by customed R script.

In addition, the h5 could be reanalyzed by Cellranger by handing the output of droputils

##### Processing scRNA-seq data for velocity analysis

Expression matrices generated above were imported to Seurat 4.1.0(Hao et al., 2021) or MUDAN 0.1.0(Purroy et al., 2018), which were first log normalized and scaled. The number of principal component analysis (PCA) was mainly determined by the elbow graph, which guided the unsupervised clustering. The number of clusters were determined by multi-resolution through clustree 0.4.4(Zappia & Oshlack, 2018). The clustering results were mainly displayed and analyzed by tSNE and UMAP. Cell populations were mainly determined by marker genes (Spermary cells see supplementary note table S1, limbal cells see supplementary table S2).

##### Theory of Region velocity and length velocity

The model is shown in Fig. 1a. The theory of length velocity is almost the same as RNA velocity and the theory of Region velocity is inspired and inferred by RNA velocity. Region velocity includes steady-state model and dynamical model using EM algorithm. The detailed inference and computational framework are elaborated in supplementary notes. Simulation of exons and introns using Region velocity is completed by computation framework of EM algorithm excluded iteration step (step 4 in supplementary notes) to solve the switch time from transcription process to splicing and degradation process. With switch time, equation 11 and 12 in supplementary notes using parameters from steady-state model are used to simulate the expected exons and introns counts.

#### Velocity analysis pipeline

RNA velocity is implemented in R package (velocyto.R)(La Manno et al., 2018) of original framework. All greedy balanced KNN algorithm in two samples used the default parameters. The velocity is estimated using gene.relative.velocity.estimates function with parameters ‘fit.quantile = 0.05, min.nmat.emat.correlation = 0.2, min.nmat.emat.slope = 0.2, kCells = 10’ from expression matrices of unspliced and spliced counts. The projection plot is drawn using show.velocity.on.embedding.cor function with parameters ‘show.grid.flow = TRUE’. Pipeline of length velocity is similar as that of RNA velocity except the input matrices from total length of unspliced and spliced mRNA. Region velocity is implemented in new R package – Regionvelocity (https://github.com/Dekayzc/Regionvelocity) which contained steady state model and dynamics model from matrices of exons and introns counts. Detailed pipelines can be found in protocol.io.

##### Acknowledgment

We sincerely thank Dr. Chuanle Xiao/Dr. Fei Sun for sharing the single cell data.

## Author contributions

CT designed and supervised the experiments. ZCC perform the lab experiments; CZ, WTC and YTF perform the bioinformatics data analysis. All authors combinedly performed the data analysis. All authors have read and approved the final manuscript draft.

---

**Extended Data Fig.1.**
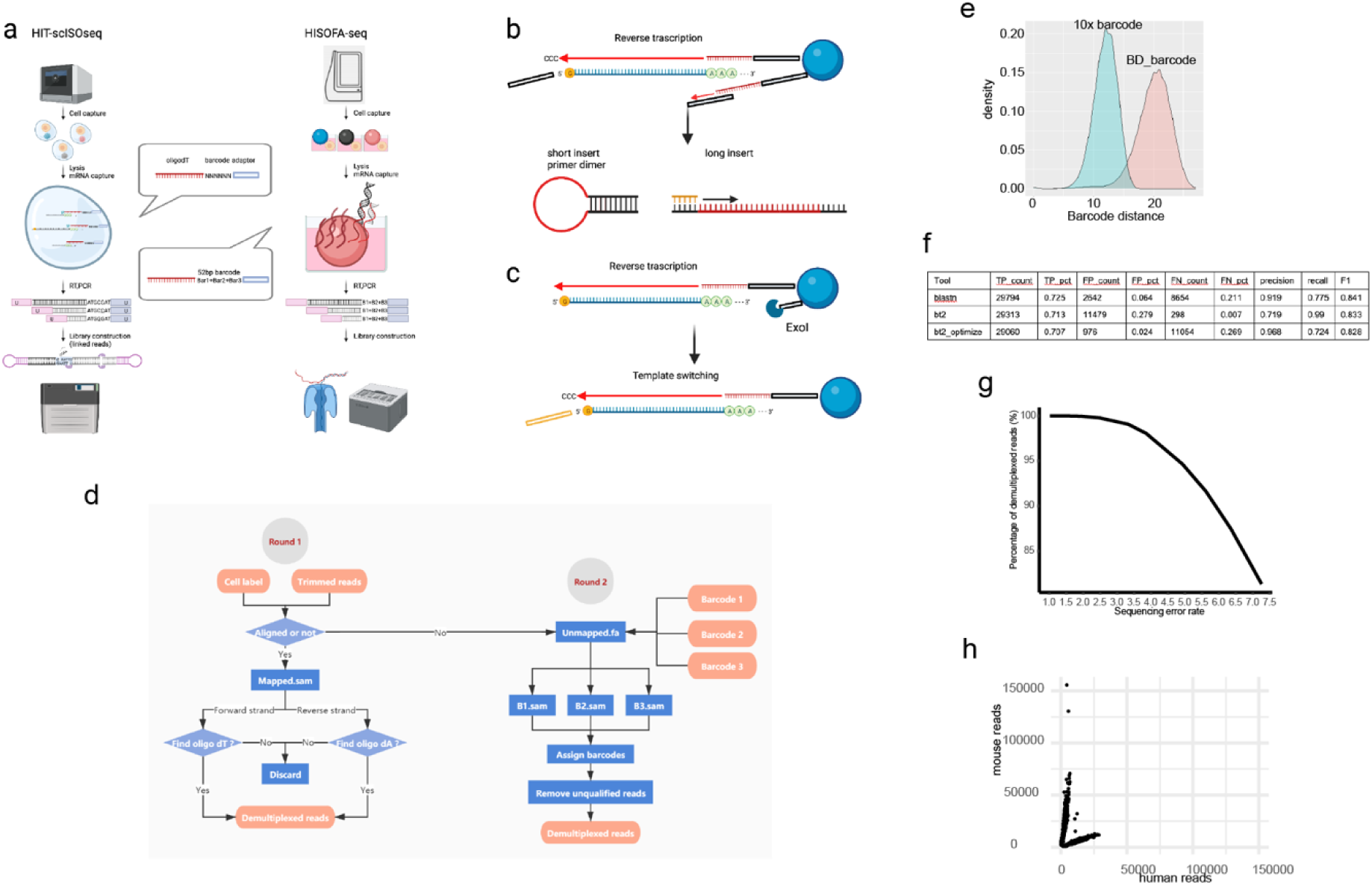
The principle of the HIT-scisoseq and HISOFA-seq. (a) In our previous research ^1^, we developed the first 20million reads full-length single cell platform HIT-scisoseq, by using the 10x Genomics Chromium Controller (10xGCC) and Pacbio sequel II. The cells were captured in droplets 1 and labeled by the 1million GEMs (Gel Bead-In EMulsions) with 9bp barcode sequence. Then the reverse transcription tagged the other adaptor on the 5’ end by the template switching activity. PCR amplification incorporated the specific uracil containing primers. The sticky end PCR fragments, generated by USER, were linked together as the head-to-tail ligation. The head-to-tail ligation produced 5-4 fold (16million) than the conventional Pacbio sequel II (4million). The more details could be found on our previous research^1^. In this research, we planned to take advantage of the nanopore high-throughput sequencing to generate the 100 million cDNA reads at low cost. We developed another customer accessible platform HISOFA-seq, combining the BD Rhapsody system and the nanopore sequencing. The single cells were captured in microarray and labeled by beads with 52bp mega barcodes, the large Levenshtein distance among which literately could compensate for the accuracy deficiency (70%∼90% accuracy) of nanopore sequencing. In the following reverse transcription, our previous HIT-scISOseq and other researches^2-5^ tagged the same sequence on the 5’ end, inhibiting the dimmer/short fragments (<500bp) amplification with the stem-loop structure(b).(c) To sequence the full-scale RNAs (nascent, stable, short degrading RNAs) and avoid the dimmer amplification, we used the exonuclease I (ExoI) to remove excess primers on beads after reverse transcription and then tagged the other 5’ adaptor on recycled beads. The PCR efficiently amplified the full-scale cDNAs, offering the holographic view of the natural transcriptomic landscape. (d) Schematic representation of the barcode demultiplexing process. A two-step pipeline was designed for BD barcodes sequenced with Oxford Nanopore platform. The first step is done by aligning reads to the BD cell label whitelist. The result was then validated by the presence of oligo dT in forward aligned reads and oligo dA in reverse aligned reads. Any unaligned reads were collected and aligned to three barcodes separately. Only reads aligned to all three barcodes were regarded as successfully recovered reads(2^ND^ round is optional for many users). (e) The density plot of BD barcode and 10X barcode edit distances. BD_barocode (HSOFA-seq) have 2x larger distance than the 10x barcode system. (f) Performance of the first round of demultiplexing with different algorithms. HISOFA-seq were based on the bt2_optimized algorithm, which have the comparable specificity and sensitivity with blastn. (g) Simulated data showing the percentage of demultiplexed reads while increasing sequencing error rate. 10,000 reads with a variety of error rates were simulated with ART software followed by out demultiplexing pipeline. Percentage of recovered breads were calculated for different error rates. (h) Dot plot of human-mouse collision. A mixture of human and mouse cells was used to estimate barcode collision.

**Extend Data Fig.2.**
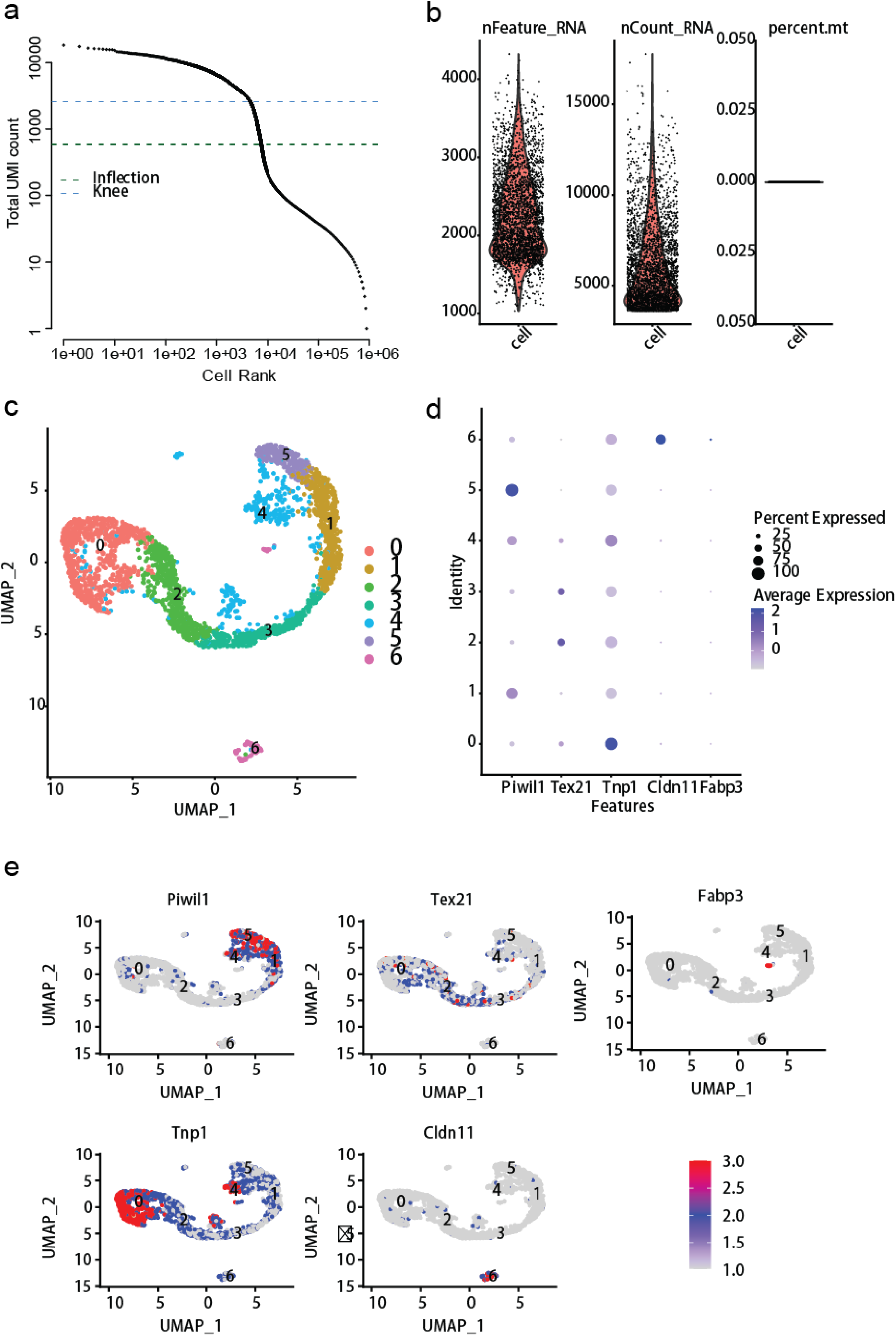
The performance of HISOFA-seq from mouse spermary cells. (a) Elbow plot to determine cell numbers. (b) The density of gene numbers and gene counts in each cell. Percent.mt represents the percentage of mitochondrial genes. (c) Cell clusters onto UMAP plot. (n=3,001 cells). (d) Dot plot of marker genes-*Piwil1* (spermatocytes), *Tex21* (round spermatids), *Tnp1* (elongating spermatids), *Cldn11* (Sertoli cells) and *Fabp3* (Leydig cells). (e) Feature plots onto UMAP of marker genes.

**Supplementary figure 3.**
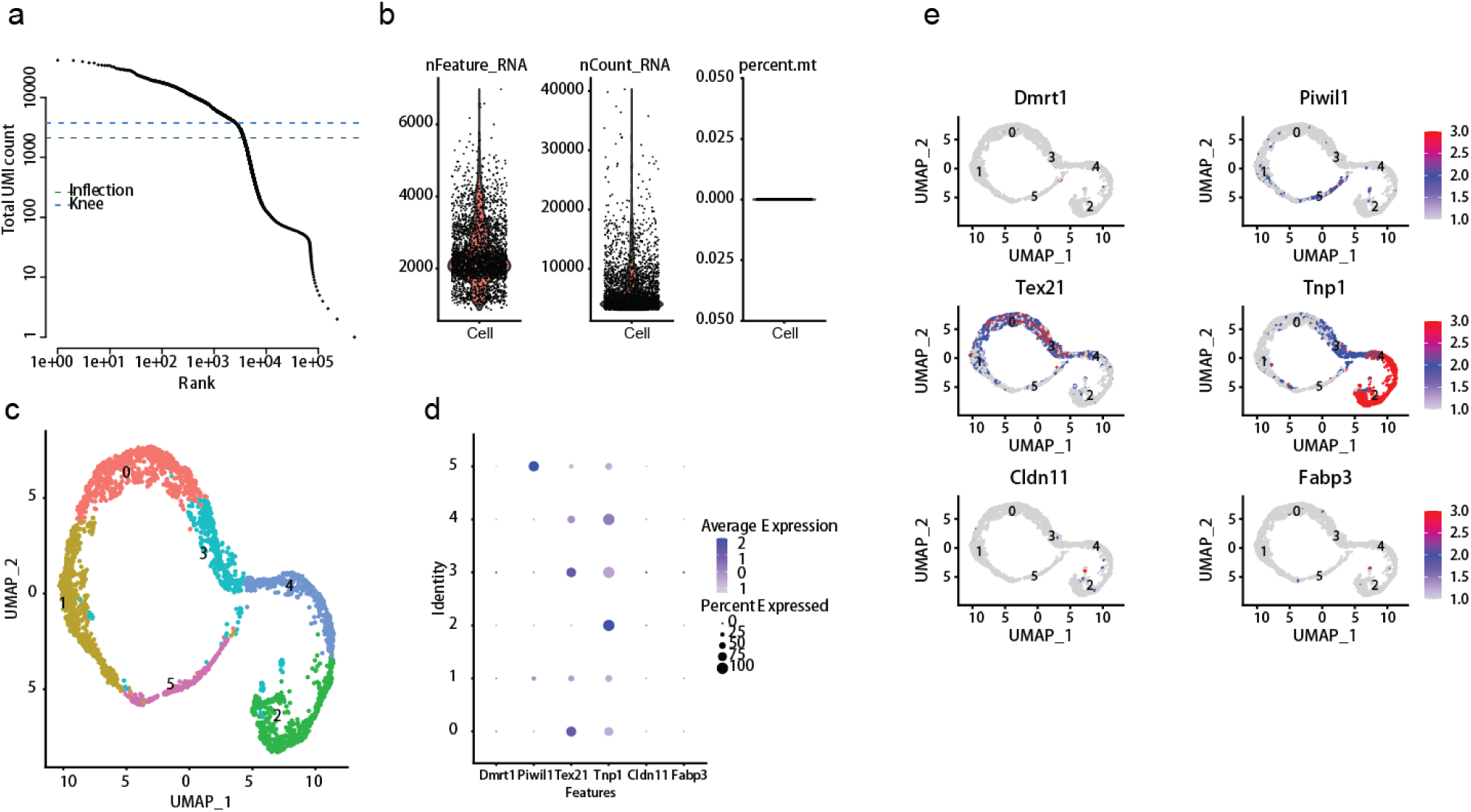
The performance of HIT-scISOseq from mouse spermary cells. (a) Elbow plot to determine cell numbers. (b) The density of gene numbers and gene counts in each cell. Percent.mt represents the percentage of mitochondrial genes. (c) Cell clusters onto UMAP plot. (n=3,001 cells). (d) Dot plot of marker genes-*Dmrt1* (spermatogonia), *Piwil1* (spermatocytes), *Tex21* (round spermatids), *Tnp1* (elongating spermatids), *Cldn11* (Sertoli cells) and *Fabp3* (Leydig cells). (e) Feature plots onto UMAP of marker genes.

**Extended Data Fig.4.**
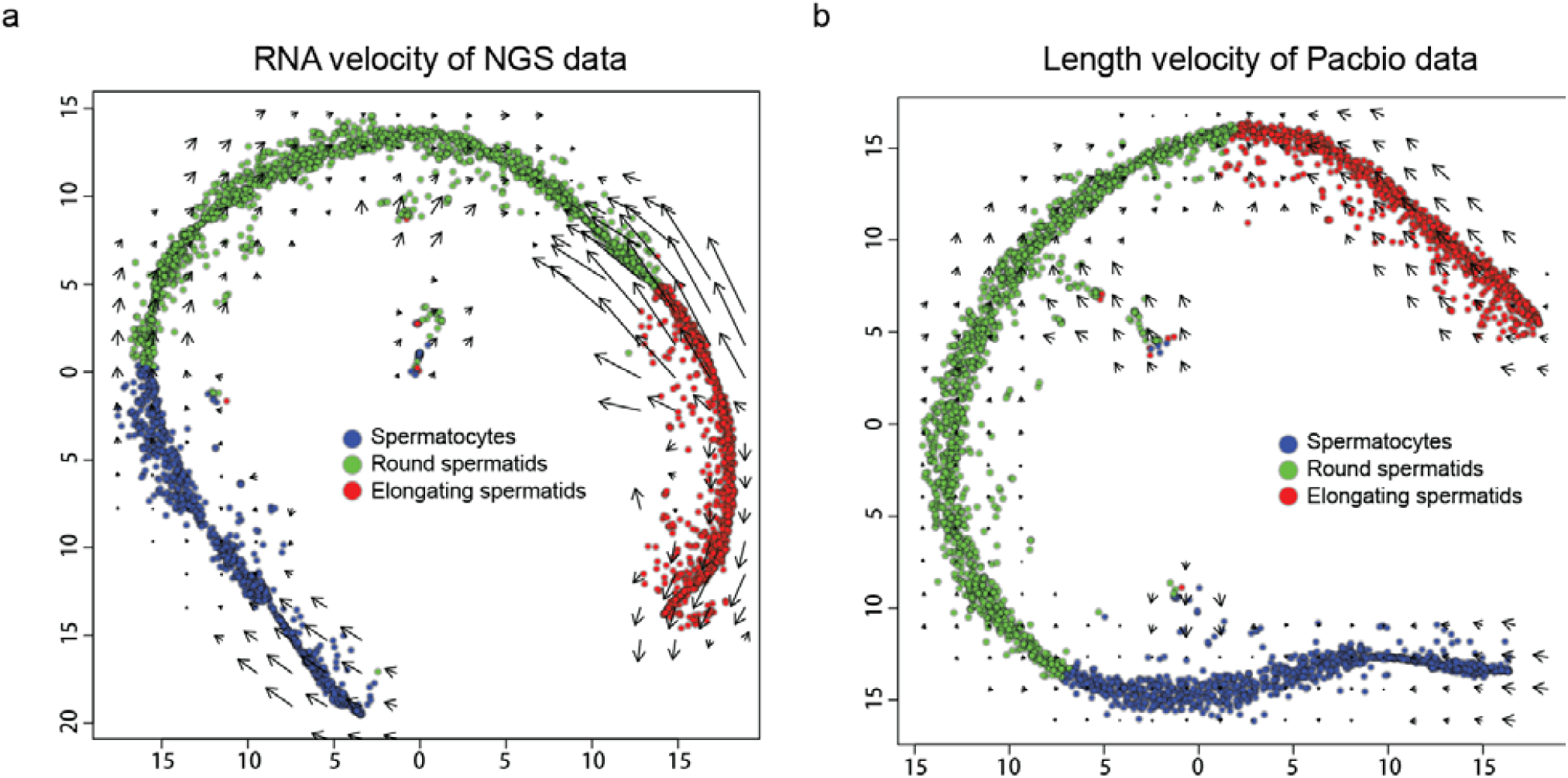
Velocity filed projected onto tSNE plot of RNA velocity and length velocity from scRNA data of the same mouse spermary cells (n=3,001 cells) using (a) 10x NGS. Arrows means the average speed on a defined grid (number of grids=20). Blue points represent spermatocytes, green points represent round spermatids and red points represent elongating spermatids. Cell populations are also determined by same marker genes as Extended Data Fig. 3. (b) HIT-scISOseq. The length for one gene in a cell is calculated as total length of all counts of a gene which are divided to spliced counts and unspliced counts in advance.

**Extend Data Fig.5.**
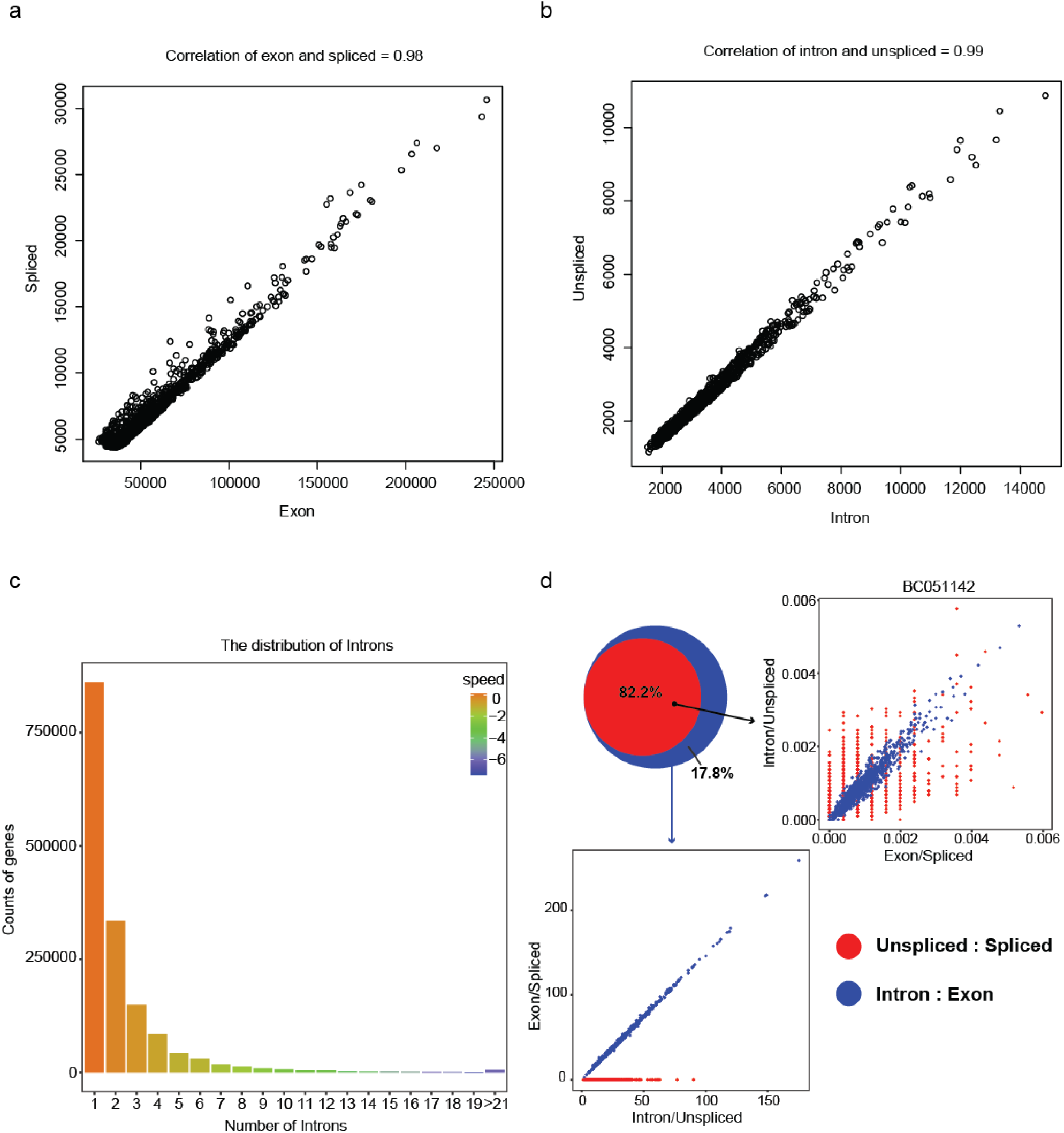
Relationship among introns, unspliced counts, exons and spliced counts. (a) The correlation between Exons and spliced counts. (b) The correlation between introns and unspliced counts. (c) The distribution of introns corresponding to their speed of introns change. The speed above 0 means increased trend of introns and below 0 means decreased trend of introns. (d) Comparison between correlation of unspliced counts and spliced counts and that of introns and exons. Top left insert is Venn plot of two correlation profiles. Top right insert is the scatter plot of a specific gene in different cells existed in both correlation profiles. Bottom left insert is the scatter plot of all gene in different cells only existed in correlation of exons and introns.

**Extended Data Fig.6.**
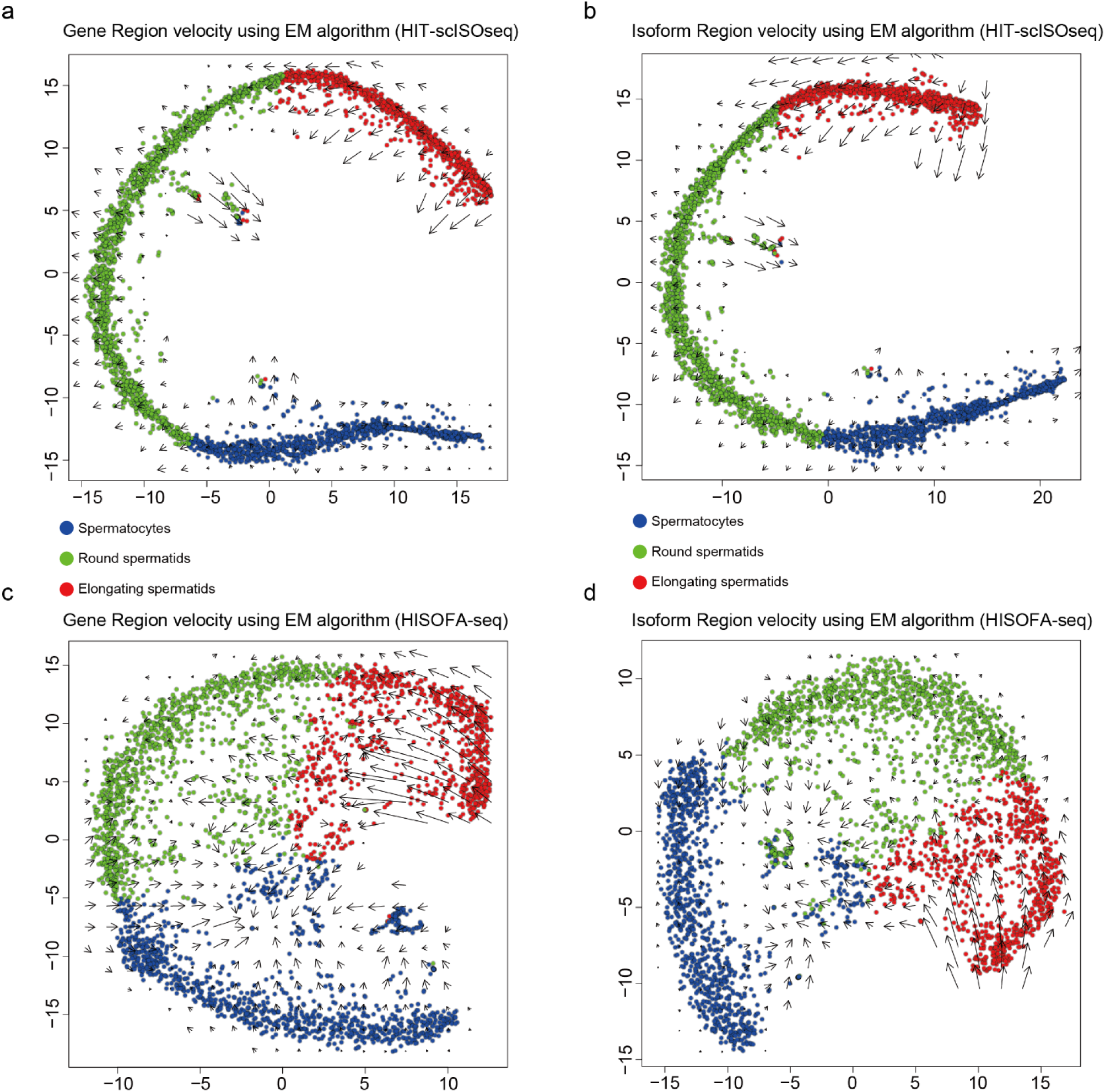
Velocity filed projected onto tSNE plot of Region velocity with dynamics model (EM algorithm) from full-scale scRNA data of the mouse spermary cells (n=3,001 cells) using (a) HIT-scISOseq in gene level. Arrows means the average speed on a defined grid (number of grids=20). Blue points represent spermatocytes, green points represent round spermatids and red points represent elongating spermatids. Cell populations are identified with marker genes (Extended Data Fig. 3). (b) HIT-scISOseq in isoform level in which reads are clustered to isoforms. (c) HISOFA-seq in gene level. Cell populations are identified with same marker genes (Extended Data Fig. 2). (d) HISOFA-seq in isoform level.

**Extended Data Fig. 7.**
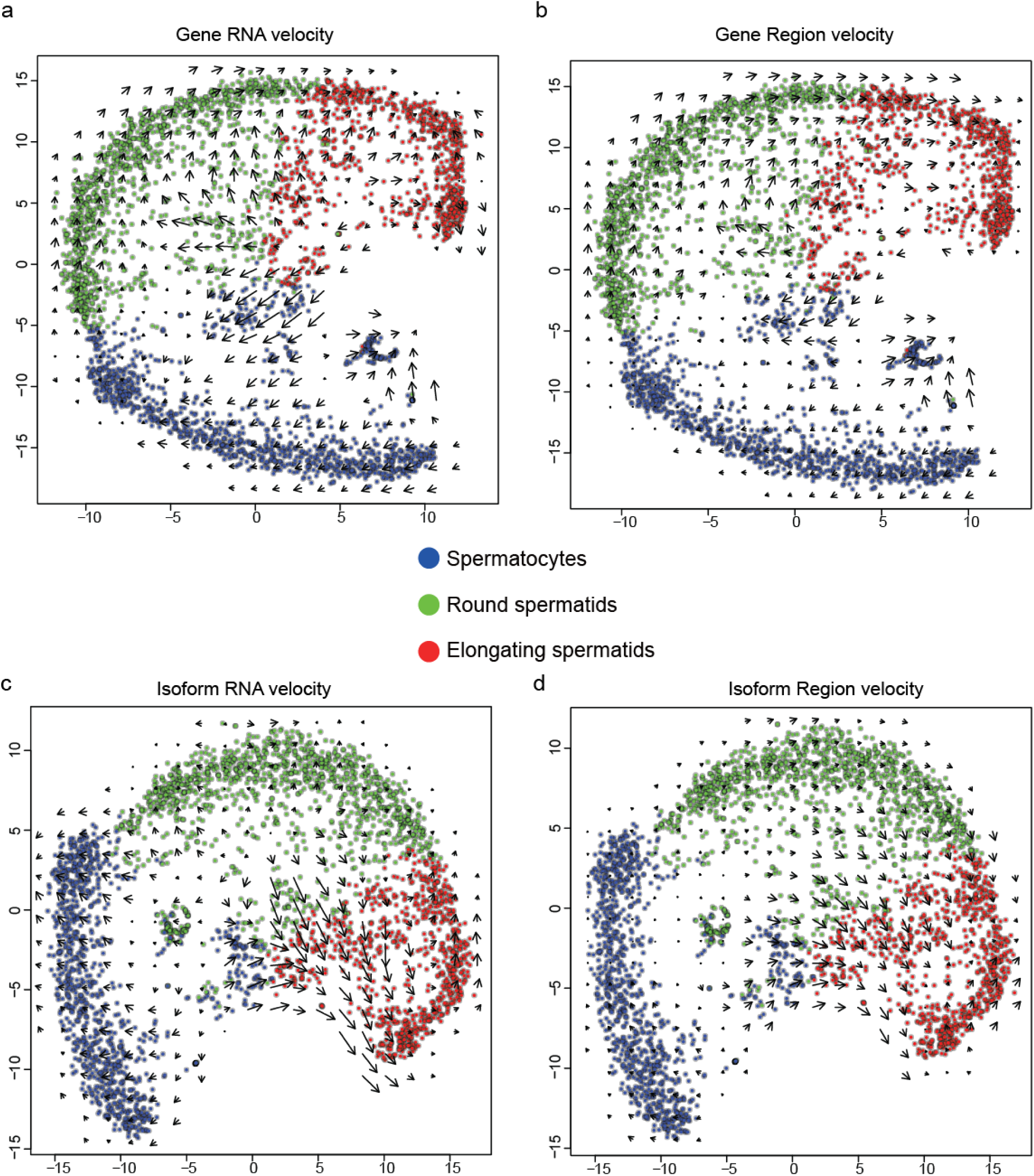
Velocity filed projected onto tSNE plot from full-scale scRNA data of the mouse spermary cells (n=3,001 cells) using HISOFA-seq with (a) RNA velocity model in gene level using spliced counts and unspliced counts as observations. Arrows means the average speed on a defined grid (number of grids=20). Gene level means the gene counts are determined by original alignment to reference without isoform calling. Blue points represent spermatocytes, green points represent round spermatids and red points represent elongating spermatids. Cell populations are identified with marker genes (Extended Data Fig. 2). (b) Region velocity model in gene level using exons and introns as observations. (c) RNA velocity model in isoform level in which reads are clustered to isoforms. The isoform clustering method might reduce the influcence of error rate of ONT sequencing. The gene counts are determined by the sum of all isoforms belonged to the gene. The main difference between gene level and isoform levels are the difference of gene counts method. (d) Region velocity model in isoform level.

**Extended Data Fig. 8.**
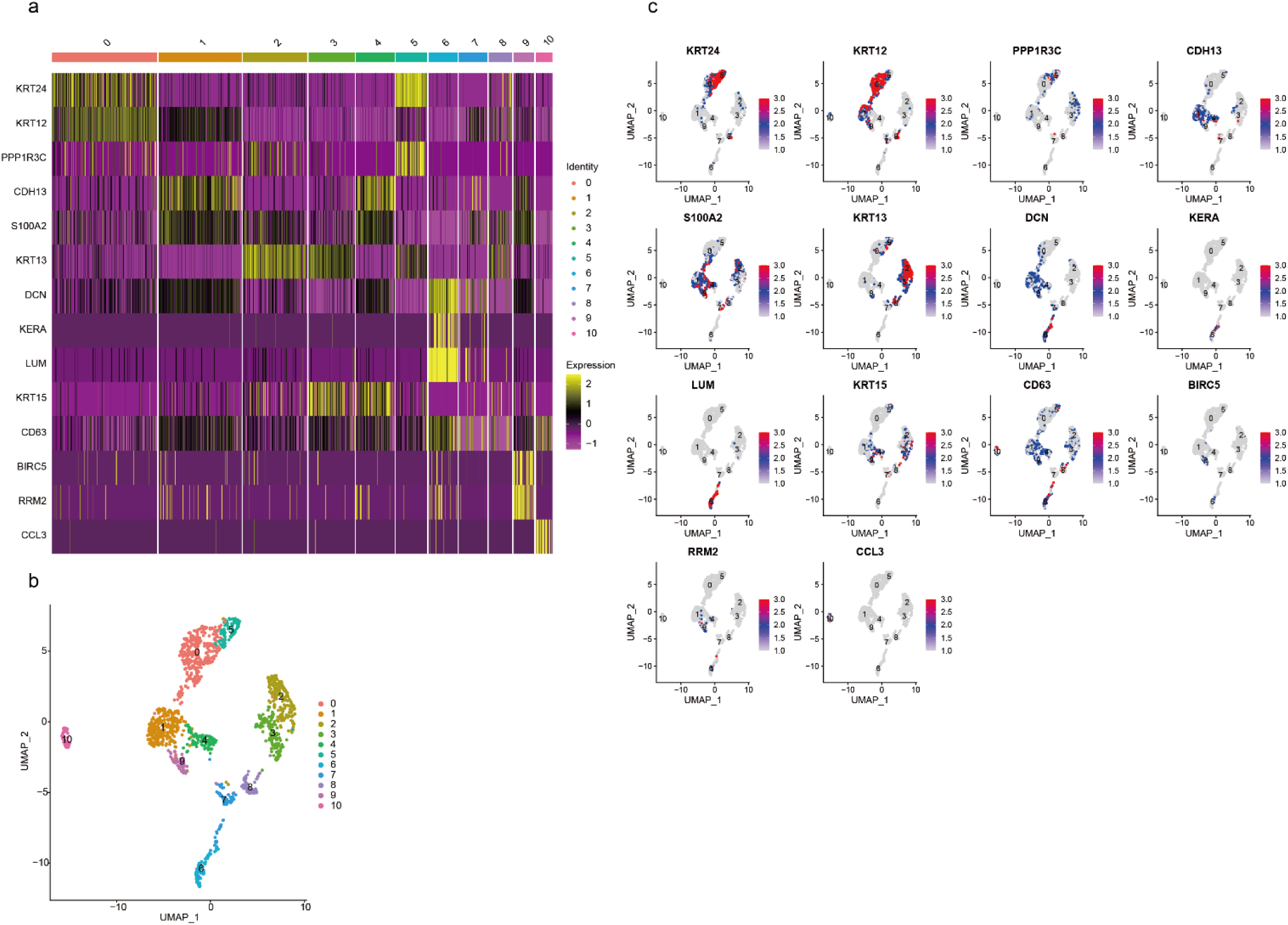
Identification of cell population for cynomolgus monkeys’ limbal cells (n=1,417 cells) using HIT-scISOseq. (a) Heatmap of marker genes including *KRT24, KRT12, PPP1R3C* (Corneal epithelial cells), *CDH13, S100A2* (Limbal progenitor cells), *KRT13* (Conjunctival epithelium), *DCN, KERA, LUM* (Corneal stromal cells), *KRT15, CD63* (Limbal stem cells), *BIRC5, RRM2* (Amplifying cells), *CCL3* (Immune cells). Cluster 0,5 expressed high in gene *KRT24, KRT12, PPP1R3C*, indicating them as corneal epithelial cells. Cluster 1,4 expressed high in gene *CDH13, S100A2*, indicating them as limbal progenitor cells. Cluster 2,3 expressed high in gene *KRT13*, indicating them as conjunctival epithelium. Cluster 6 expressed high in gene *DCN, KERA, LUM*, indicating it as corneal stromal cells. Cluster 7,8 expressed high in serveral limbal cells related genes including *KRT15, CD63*, indicating their potential as stem cells. Cluster 9 expressed high in limbal cells related gene especially related with limbal progenitor and limbal stem cells. However, cluster 9 showed a high expression in gene *BIRC5, RRM2*, indicating amplification process existed in cluster 9. Therefore, cluster 9 showed potential as transient amplifying cells. Cluster 10 expressed high in gene *CCL3*, indicating it as immune cells. (b) Distribution of clusters in UMAP plot. (c) Dot plot of marker genes.

