## supplementary notes for "Improving the RNA velocity approach using long-read single cell sequencing"

**Region velocity resolve the long-read RNA velocity in single cells**

### Supplementary notes: Mathematical inference and computational methods.

#### Inference of Length velocity

One of the highlights of long-read scRNA-seq technology (HIT-scISOseq/HISOFA-seq in main content) is the detection of full-scale mRNAs (from nascent to degrading RNAs), of which the length is suitable for investigating its model like RNA velocity. According to RNA transcription dynamics model (Fig. 1a), the length of mRNA also varies over time in the process of transcription, splicing and degradation. The basic reaction kinetics can be described as follow by each gene independently

$$\frac{dl}{dt} = \alpha - \beta \cdot \%u \cdot l(t) - \gamma \cdot \%s \cdot l(t) \qquad (1)$$

where, like RNA velocity model, we also consider transcription rate ($\alpha$), splicing rate ($\beta$) and degradation rate ($\gamma$) as time-independent. The length of mRNA is reduced in both splicing and degradation process partially ($\%u$ and $\%s$, respectively). In fact, $\%u$ and $\%s$ can be regarded as the proportion of unspliced and spliced mRNA's length. Therefore, equation 1 can be decomposed to two following equations:

$$\frac{dl_u}{dt} = \alpha - \beta l_u(t) \qquad (2)$$

$$\frac{dl_s}{dt} = \beta l_u(t) - \gamma l_s(t). \qquad (3)$$

Obviously, equation 2 and 3 is similar as the equations of original RNA velocity[1], which substitute u(t) as $l_u$(t) and s(t) as $l_s$(t). So, the computational framework of RNA velocity can directly apply to that of length velocity. The only difference between two computational frameworks is the input files which are mRNA counts and mRNA length respectively. However, the length of mRNA has two types. One is the total length for a gene in a cell, and the other one is the average length. Using total length could include the influence of mRNA abundance, which will be more consistent with RNA velocity. Therefore, the result of length velocity in this study is similar as RNA velocity since we use total length as observation values to predict cell fate. Practically, the average length could highlight the influence of length change over cell fate. Hence, the computational framework in this study provides two kinds of length as input to RNA velocity package[1].

**Inference of Region velocity**

Besides length, long-read scRNA-seq technology can survey substantial observations, which have potential to predict the cell fate. The observation of exon-intron plots inspires the idea of Region velocity. As depicted in main content, the variation number of introns and exons in mRNA is different from that of unspliced and spliced counts in the process of splicing and degradation. By the law of mass action, the rate equations for each gene independently of which the expected exons count *e* and introns count *i* evolve over time can be simply described as follow:

$$\frac{di}{dt} = \alpha_i(t) - \beta i(t), \qquad (4)$$

$$\frac{de}{dt} = \alpha_e(t) - \gamma e(t), \qquad (5)$$

where, $\alpha_i$(t) and $\alpha_e$(t) are the transcription rate of *i* and *e* respectively. However, $\alpha_i$(t) and $\alpha_e$(t) have specific relations as the nascent mRNA produced during transcription process are consist of introns and exons. So, we introduce a parameter - ratio of introns in nascent mRNA ($\theta$) as this parameter can be estimated more easily from observed data than $\alpha_i$(t) and $\alpha_e$(t) directly. Then equation 4 and 5 can be simplified as follow:

$$\frac{di}{dt} = \theta\alpha(t) - \beta i(t), \quad (6)$$

$$\frac{de}{dt} = (1-\theta)\alpha(t) - \gamma e(t), \quad (7)$$

where, splicing rate ($\beta$) and degradation rate ($\gamma$) are also considered as time-independent and transcription rate ($\alpha$) is conditionally constant as follow:

$$\alpha(t) = \begin{cases} \alpha & t \le t_s, \\ 0 & t > t_s, \end{cases} \quad (8)$$

where, $t_s$ is the switch time of the transcription process, which means that when time is smaller than $t_s$, the transcription process would be continuously operated and the transcription rate ($\alpha$) would be time-independent value $\alpha$, and when time is larger than $t_s$, the transcription process would be terminated and $\alpha$ would be 0. The assumption of Eq. 8 would ensure the consistent trend of variation of exons and introns over time. The analytical solution to the first order linear differential equations (Eq. 6 and 7) with assumption $\alpha$(t)= $\alpha$ can be

$$i(t) = i_0 e^{-\beta t} + \frac{\theta\alpha}{\beta}(1 - e^{-\beta t}), \quad (9)$$

$$e(t) = e_0 e^{-\gamma t} + \frac{(1-\theta)\alpha}{\gamma}(1 - e^{-\gamma t}), \quad (10)$$

where $i(0)=i_0$, $e(0)=e_0$. Normally, we can assume $i_0=0$ and $e_0=0$. Combined with equation 8, the analytical solution can be

$$i(t) = \begin{cases} \frac{\theta\alpha}{\beta}\left(1 - e^{-\beta t}\right) & t \le t_s \\ i_s e^{-\beta(t-t_s)} \qquad i_s = \frac{\theta\alpha}{\beta}\left(1 - e^{-\beta t_s}\right), & t > t_s \end{cases} \quad (11)$$

$$e(t)=\begin{cases}\dfrac{(1-\theta)\alpha}{\gamma}(1-e^{-\gamma t}) & t\le t_s\\ e_s e^{-\gamma(t-t_s)} \qquad e_s=\dfrac{(1-\theta)\alpha}{\gamma}(1-e^{-\gamma t_s}), & t>t_s\end{cases} \tag{12}$$

Here, equation 11 and 12 exclude the parameters $i_0$ and $e_0$ and the constant parameters that have to be inferred are $\varphi$ ($\alpha$,$\beta$,$\gamma$,$\theta$,$t_s$). Next, we will elaborate how these parameters are estimated.

**Inference of parameters and extrapolation of cell state from data using Region velocity**

**Steady-state model**

As depicted in main content (Fig. 1e), *e* and *i* showed the trend of steady-state model according to original RNA velocity framework. Therefore, to infer the parameters from data, the idea of steady-state is borrowed[1]. In steady-state cells, the reduction of *e* and *i* would also be the same as the increase of *e* and *i*, which means *de/dt*=0 and *di/dt*=0. So, equation 6 and 7 would be

$$\theta\alpha(t)-\beta i(t)=0\rightarrow\ \alpha(t)=\frac{\beta i(t)}{\theta}, \tag{13}$$

$$(1-\theta)\alpha(t)-\gamma e(t)=0\rightarrow\alpha(t)=\frac{\gamma e(t)}{1-\theta}, \tag{14}$$

where $\alpha(t)$ in equation 13 should be equal to $\alpha(t)$ in equation 14. Therefore, combined with equation 13 and equation 14, we can obtain

$$i(t)=\frac{\theta\gamma e(t)}{(1-\theta)\beta}. \tag{15}$$

Let $\gamma^*=\frac{\theta\gamma}{(1-\theta)\beta}$, like steady-state model in RNA velocity framework, linear fit model using a least square fit of *e* and *i* from observed data can solve the parameter, i.e., $i\sim\gamma^* * e$.

Additionally, $\theta$ can be estimated from observed data. We know $\theta = \frac{i}{i+e}$ for a nascent gene and in a nascent gene the relationship between *e* and *i* could be $e = i + 1$. Also, the number of *e* and *i* should be maximal in the nascent state for a gene as in later splicing and degradation process, they would be reduced. Therefore, the estimation of $\theta$ from observed data can be

$$\theta = max\left(\frac{max(i)}{2max(i)+1}, \frac{max(e)-1}{2max(e)-1}\right). \quad (16)$$

With estimation of $\theta$, $\frac{\gamma}{\beta}$ can be solved from $\gamma$*. Although only estimation of $\theta$ and $\frac{\gamma}{\beta}$ in steady-state model are figured out, the extrapolation of cell state could be still completed. Current cell state could be represented by the observed value *e* of all genes. So, projected cell state could be represented by the extrapolated value *e* of all genes. For a single gene in each cell, the observed value *e* can be regarded as *e*(t) in a specific time point t. We assume the extrapolated value *e* would be obtained after Δt. Then its time point would be t+Δt and extrapolated value *e* could be *e*(t+Δt). So, based on equation 10, *e*(Δt) = *e*(t+Δt) - *e*(t) could be

$$e(\Delta t) = (e^{-\gamma\Delta t} - 1)(e_0 e^{-\gamma t} - \frac{(1-\theta)\alpha}{\gamma} e^{-\gamma t}), \quad (17)$$

where we can figure out $e_0 e^{-\gamma t} - \frac{(1-\theta)\alpha}{\gamma} e^{-\gamma t} = e(t) - \frac{(1-\theta)\alpha}{\gamma}$ according to equation 10 and $\alpha = \frac{\beta i(t)}{\theta}$ according to equation 13. Then, equation 17 can be simplified as follow:

$$e(\Delta t) = (e^{-\gamma\Delta t} - 1)\left(e(t) - \frac{(1-\theta)i(t)}{\theta\frac{\gamma}{\beta}}\right). \quad (18)$$

So, the extrapolated value e(t+Δt) can be estimated from estimated parameters $\theta$ and $\frac{\gamma}{\beta}$ and observed value *e* and *i* with a given period Δt. Except parameters inference and extrapolation equation building, other preprocessing, balance KNN pooling, visualization of velocity, linear regression weights and top/bottom quantiles definition are referred from RNA velocity framework to avoid reinventing wheels[1].

**Dynamical model**

As steady-state model only completed estimation of two parameters and Volker Bergen et al. indicated the defect of steady-state model[2], we continue using dynamical model to infer the parameters $\varphi$ ($\alpha$,$\beta$,$\gamma$,$\theta$,$t_s$). After studying equation 11 and 12, we find we can only obtain observed value of *e* and *i*. and another variable time t cannot be observed. Naturally, the hidden variable question drives us to consider Expectation–maximization (EM) algorithm[3].

Normally, the first step of EM algorithm is initialization of parameters $\varphi$. As we have obtained estimation of part parameters $\varphi$ from steady-state model, initialization of parameters $\varphi$ could be completed based on steady-state model. However, as time is a hidden variable, switch time $t_s$ should also be a hidden variable. So, the parameters $\varphi$ to be initialized would be reduced to $\varphi$' ($\alpha$,$\beta$,$\gamma$,$\theta$). Like RNA velocity[1], we can set $\beta$=1. Then $\gamma$ can be solved from linear fit result $\gamma$*. So, only $\alpha$ is not assigned an initialed value. According to equation 13, $\alpha$ can be solve from observed value *i*. Here we set $\alpha = \beta/\theta * mean(i)$. As parameters $\varphi$' are initialized, the second step of EM algorithm is expectation step of hidden variables.

Like RNA velocity[2], we can assume the difference between observed value and expected value is normal distribution in which mean value is 0 and variance is σ2. Then the likelihood for a gene can be

$$L(\varphi') = \frac{1}{2\pi\sigma^2} exp\left(\frac{1}{n}\sum_{c}^{n} - \frac{(e_c - e(t_c, \varphi'))^2 + (i_c - i(t_c, \varphi'))^2}{2\sigma^2}\right), \tag{19}$$

where $e(t_c, \varphi')$ and $i(t_c, \varphi')$ are the solution of equation 11 and 12 at time $t_c$, $t_s$ and parameters $\varphi'$ for the cell *c*. Then the log-likelihood $l(\varphi')$ have this proportionality:

$$l(\varphi') \propto -\sum_{i}^{n} (e_c - e(t_c, \varphi'))^2 + (i_c - i(t_c, \varphi'))^2. \tag{20}$$

So, the hidden variable *t* for different cell *c* can be solved by maximum likelihood estimation (MLE) of $l(\varphi')$:

$$t_c = \underset{\varphi'}{\operatorname{argmax}}\, l(\varphi') = \underset{\varphi'}{\operatorname{argmin}}((e_c - e(t_c, \varphi'))^2 + (i_c - i(t_c, \varphi'))^2). \tag{21}$$

As the derivative method will encounter the transcendental equation, in this study, $t_c$ is solved by quasi-Newton method which specifically refers to Broyden–Fletcher–Goldfarb–Shanno (BFGS) algorithm[4] with low memory in R package[5]. However, only part of $t_c$ can be solved since a key hidden variable switch time $t_s$ is still unknown. To solve $t_s$, we should give the criteria when $t \le t_s$ and $t > t_s$. Switch time is defined as the time gene changed from transcription process to splicing and degradation process. In other words, as for equation 6 and 7, in transcription process, we have $di/dt > 0$ and $de/dt > 0$. Observing equation 11 and 12, when $t \le t_s$, *e*(t) and *i*(t) can be solved without hidden variable $t_s$. Hence, the hidden variable *t* in cells of transcription process can be solved based on equation 21. Additionally, the condition $di/dt > 0$ and $de/dt > 0$ can be simplified to $i < \theta * \alpha/\beta$ and $e < (1-\theta) * \alpha/\gamma$. So, $t_s$ can be solved by

$$t_s = max(t_c) \quad c \in (i_c < \frac{\theta\alpha}{\beta}, e_c < \frac{(1-\theta)\alpha}{\gamma}). \qquad (22)$$

Then, time *t* of other cells that do not belong to cells in transcription process will be solved by equation 21 with $t_S$. So, in this expectation step, hidden variable *h*(*t*,*t*s) is estimated and the third step of EM algorithm maximization step is available.

The hidden variable *h*(*t*,*t*s) is able to re-estimate parameters *φ'*. Similar as equation 21, parameters *φ'* for one gene can be solved by observed values in all cells as follow:

$$\varphi' = \underset{t}{\operatorname{argmax}}\, l(\varphi') = \underset{t}{\operatorname{argmin}} \sum_{c}^{n} ((e_c - e(t_c, \varphi'))^2 + (i_c - i(t_c, \varphi'))^2). \qquad (23)$$

Parameters *φ'* are still solved by BFGS. The final step of EM algorithm is the iterations of second step and third till convergence. The convergence condition in this study is set to 100 (default but user-defined) iterations or $(\varphi'_{iteration+1} - \varphi'_{iteration}) / \varphi'_{iteration} < 1\%$. In summary, the EM algorithm in this study can be stated as follow:

Step 1. Initialization of parameters *φ*' (*α*,*β*,*γ*,*θ*) from stead-state model.

Step 2. Inference of hidden variable *h*(*t*,*t*s) for each gene from current parameters *φ*'.

Step 3. Inference of parameters *φ*' (*α*,*β*,*γ*,*θ*) for each gene from *h*(*t*,*t*s) in step 2.

Step 4. Iterations of step 2 and step3 till convergence.

The extrapolation of cell state is easier as all parameters are estimated. For a specific period Δt, the projected e(t+Δt) can be solved with equation 11 and 12 for a single gene. Then overall velocity of a cell is comprised of all genes in a cell. The cell fate flow can be observed from current cell point to projected cell point in embedded tSNE or UMAP plot. Visualization of velocity in dynamical model are still referred from RNA velocity framework.

1. La Manno, G. et al. RNA velocity of single cells. *Nature* **560**, 494-498 (2018).
2. Bergen, V., Lange, M., Peidli, S., Wolf, F.A. & Theis, F.J. Generalizing RNA velocity to transient cell states through dynamical modeling. *Nature biotechnology* **38**, 1408-1414 (2020).
3. Dempster, A.P., Laird, N.M. & Rubin, D.B.J.J.o.t.R.S.S.S.B. Maximum likelihood from incomplete data via the EM algorithm. **39**, 1-22 (1977).
4. Byrd, R.H., Lu, P., Nocedal, J. & Zhu, C.J.S.J.o.s.c. A limited memory algorithm for bound constrained optimization. **16**, 1190-1208 (1995).
5. Computing, R.J.V.R.C.T. R: A language and environment for statistical computing. (2013).
